# Development and simulation of fully glycosylated molecular models of ACE2-Fc fusion proteins and their interaction with the SARS-CoV-2 spike protein binding domain

**DOI:** 10.1101/2020.05.05.079558

**Authors:** Austen Bernardi, Yihan Huang, Bradley Harris, Yongao Xiong, Somen Nandi, Karen A. McDonald, Roland Faller

## Abstract

We develop fully glycosylated computational models of ACE2-Fc fusion proteins which are promising targets for a COVID-19 therapeutic. These models are tested in their interaction with a fragment of the receptor-binding domain (RBD) of the Spike Protein S of the SARS-CoV-2 virus, via atomistic molecular dynamics simulations. We see that some ACE2 glycans interact with the S fragments, and glycans are influencing the conformation of the ACE2 receptor. Additionally, we optimize algorithms for protein glycosylation modelling in order to expedite future model development. All models and algorithms are openly available.

## Introduction

As of June 29, 2020 more than 10 Million people have been confirmed to be infected with coronavirus disease 2019 (COVID-19) which is caused by the severe acute respiratory syndrome coronavirus 2 (SARS-CoV-2). This zoonotic pandemic has disrupted society worldwide on a peacetime-unprecedented scale. It also spurred a wide range of scientific endeavors to attack the various aspects of this disease. As the disease spreads there is a critical need for tools that enable the strategic design of biopharmaceutical countermeasures. We are here addressing computationally a molecular approach to aid in the design of a specific class of potential COVID-19 countermeasures.

The genomic sequence of the virus responsible for COVID-19, SARS-CoV-2, was made available in January 2020 (1), providing critical information on the primary amino acid sequences of potential targets. A particularly important target is the SARS-CoV-2 spike (S) protein that is responsible for the first step in the viral infection process, binding to human cells via the angiotensin converting enzyme 2 (hACE2) receptor. The conserved expression and interaction of ACE2 indicates a wide range of hosts (human and non-human) for SARS-CoV-2 (2). The S protein contains two domains S1 and S2 on each monomer. It is a homotrimer with each monomer comprised of 1281 amino acids. The monomers are expected to be highly glycosylated with 22 N-linked glycosylation sequons and 4 O-linked predicted glycosylation sites (3), although only 16 N-linked glycosylation sites were observed in a cryo-EM map of S produced in HEK293F cells (4). Very recently, Watanabe et al. performed site-specific glycoform analysis of full-length trimeric S protein made recombinantly in transfected HEK293F cells (5). Their analysis showed high occupancy at all 22 sites, with about 14 sites classified as complex, 2 sites as oligomannose, and the remaining sites containing mixtures of oligomannose, hybrid and complex glycan structures. Seven of the sites with complex glycoforms, including the 2 sites on the RBD, also had a high degree (>95%) of core fucosylation. Viral coat proteins are often glycosylated which helps pathogens evade the host immune system, modulate access of host proteases, and can enhance cellular attachment through modification of protein structure and/or direct participation at the viral coat protein/cell receptor interface. These glycans are, however, only partially resolved in the experimental structure such that a computational approach is helpful to predict their behavior.

The human ACE2 protein is a 788 amino acid integral membrane protein with seven N-linked glycosylation sites in the extracellular domain. The binding kinetics between the SARS-CoV-2 spike protein and the hACE2 receptor will depend on the 3D structures of both molecules and their molecular interactions which may be impacted by glycosylation (6–8), as has been observed for other glycosylated viral spike proteins and their human receptors. Knowledge of the spike protein and ACE2 amino acid sequences have led to the commercial availability of the spike protein, ACE2, and various fragments of these, with and without purification/fusion tags, produced recombinantly in various expression hosts including Human embryonic kidney cells (HEK293), insect cells, Chinese Hamster Ovary cells (CHO), and *E. coli*. While the availability of recombinant sources for S and ACE2 glycoproteins have greatly contributed to our understanding of the structure and interactions between these proteins, it is important to recognize that glycosylation of recombinant S and ACE proteins will depend on the host cell (9), the recombinant protein, as well as production (10, 11) and purification methods (12). As molecular models and molecular dynamics simulations can describe the interactions of proteins with glycans and the modulation of protein structure by glycans (13, 14) they are powerful tools to assess the significance of glycosylation on 3D structure and binding site interactions between the SARS-CoV-2 spike protein and the human ACE2 receptor, and to design novel biotherapeutics including optimizing glycosylation.

A promising strategy for the design of COVID-19 therapeutic proteins is a fusion of the extracellular domain of ACE2, the human receptor for SARS-CoV-2, with the Fc region of human immunoglobulin, IgG1, by a linker separating the two domains (15). The neutralization strategy behind ACE2-Fc is shown in Figure 1 (15). This therapeutic design is often called an immunoadhesin, a chimeric protein combining the ligand-binding region of the cell receptor with the constant domain of an immunoglobulin (16). These chimeric molecules form dimers through disulfide bonds between Fc domains; this bivalency increases the affinity for the ligand. The human ACE2 receptor has been shown to be the primary receptor that SARS-CoV-2 uses for entry into and infection of human cells (17, 18), although the binding site is distinct from the catalytic domain of ACE2. With an ACE2-Fc immunoadhesin the ACE2 portion can act as a circulating “bait or decoy” to bind the SARS-CoV-2 spike protein preventing it from entering human cells while the Fc region confers longer circulatory half-life, provides effector functions of the immune system to clear the virus, and allows simple well-established purification using Protein A affinity chromatography. Immunoadhesins are a distinct class of antivirals that can be used prophylactically as well as post-infection and differ from both vaccines and antibodies. Unlike vaccines, they are not intended to elicit an immune response to the viral infection, and unlike antibody therapies, their design is greatly simplified since once the cellular receptor for viral entry is identified the immunoadhesin can be quickly designed and produced.

**Figure 1.**
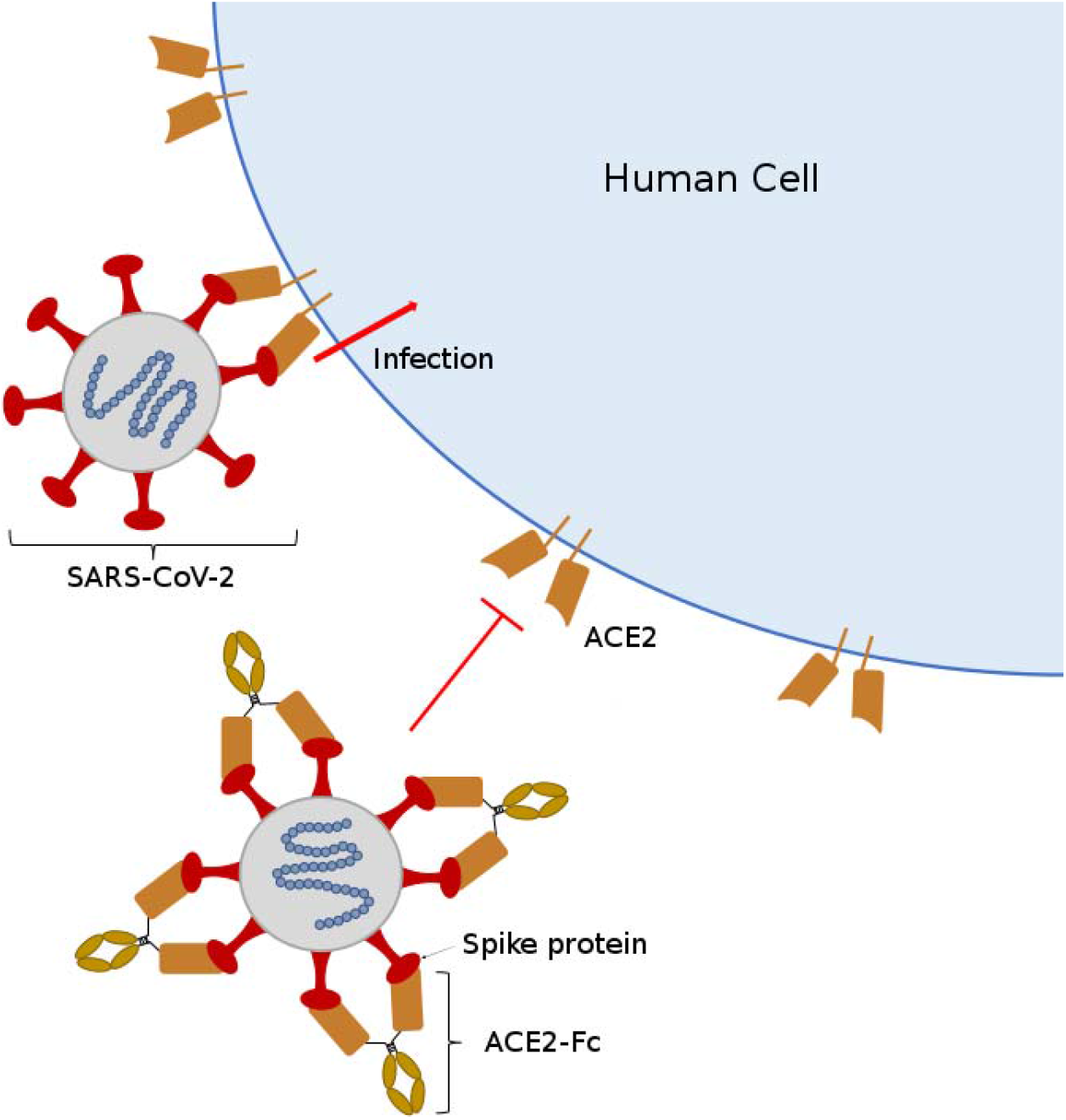
Proposed strategy for SARS-CoV-2 neutralization by ACE2-Fc immunoadhesin. ACE2-Fc binds to the spike (S) protein on the virus and blocks binding to the human cellular receptor ACE2, preventing cellular entrance of SARS-CoV-2.

This strategy also precludes the coronavirus mutating to escape binding with the ACE2-Fc protein, as it would also lose binding affinity to the native ACE2 human cell receptor resulting in a less pathogenic virus. The re-emergent SARS-CoV-1 virus in 2003-2004 had a lower affinity for human ACE2 resulting in less severe infection and no secondary transmissions (19). In this strategy the exogenous ACE2 would compensate for decreased ACE2 levels in the lungs during infection, contributing to the treatment of acute respiratory distress, and potentially reduce inflammation and reactive oxygen species in the lung (20). Most importantly, recombinant ACE2-Fc fusion proteins, with native ACE2 catalytic activity as well as a mutant version with lower ACE2 catalytic activity, produced using transfection of HEK293, have shown high affinity binding to the SARS-CoV-2 spike protein and to potently neutralize SARS-CoV-2 *in vitro* (21). Simulations are an ideal tool to optimize such a construct and guide the experimental production of ACE2-Fc.

Glycans are branched, flexible chains of carbohydrates that explore a much wider range of conformations at equilibrium conditions than the protein chain itself as the latter is typically not dynamically changing strongly from its folded form as that would affect its functionality. The faster dynamics of glycans complicates the structural and conformational characterization of glycans in laboratory experiments (22). In atomistic molecular dynamics (MD) simulations, glycan conformations can be straight-forwardly analyzed to obtain structural information, as glycan dynamics are much closer to the computational timescale than the protein dynamics. However, neighboring glycans can interact with each other and essentially lock each other in which can lead to very slow equilibration into the correct conformation (13). Therefore, algorithms are needed to generate realistic glycan configurations as glycans are regularly not fully resolved experimentally. Consequently, only a few simulations of related fully glycosylated proteins available (23–26) among them recently a proposed glycosylated model of the Spike protein (27). Very recently a short simulation of the Spike protein with glycosylation has been published which is enabling longer studies (28). Our group has made significant progress in the field of glycan modeling in recent years (13, 14, 29).

N-glycan structure is highly heterogeneous, and the relative abundance of glycans depends on the expression system for glycoprotein production. Plant-based transient expression systems are well-suited to produce recombinant ACE2-Fc under the current COVID-19 pandemic given high production speeds. Two glycovariants of ACE2-Fc are simulated in this work: one is targeted for ER retention with high mannose glycoforms, and the second is targeted for secretion with plant complex glycoforms. These glycovariants are currently being expressed and purified at UC Davis.

In order to properly understand the interaction between the spike protein and the variant ACE2 receptors bound to its fusion partner the glycosylation of both entities needs to be taken into account. The few existing computational studies of ACE2 interaction with the spike protein we are aware of are using aglycosylated proteins (30–32). Also, molecular docking studies have been performed with the older related SARS-CoV-1 virus protein implicated in the SARS epidemic in the early 2000s (33). We develop *in silico* models to predict the 3D structure of two glycosylated ACE2-Fc variants. Additionally, we evaluate the interactions between these two ACE2-Fc variants and a glycosylated spike protein fragment (SpFr) which contains the receptor binding domain of the SARS-CoV-2 spike protein.

## Materials and Methods

### Sequences and Initial Structure

ACE2-Fc is a homodimer of ACE2 bound to Fc via a synthetic linker. Two sequence variants are used in this work to model ACE2-Fc. The ACE2 and Fc domains N- and C-terminal residues for both variants, respectively, are as follows: ACE2, 18Q-740S (NCBI ID: NP_001358344.1); Fc, 109C-330K (UniProt ID: P01857). Variant 1 (Sequence Seq1 in Supporting Information; 960 amino acids) contains a C-terminal SEKDEL tag, which is used to express predominantly ER-retained proteins with high-mannose glycoforms in plant-based expression systems. Variant 2 (Sequence Seq2 in Supporting Information; 954 amino acids) does not use a C-terminal SEKDEL tag, and will express standard plant glycoforms in plant-based expression systems. Variant 2 has two mutations: H357N and H361N. These mutations are used to deactivate the standard function of ACE2, by preventing the coordination of Zn^2+^ in the active site (21). The ACE2-Fc variants contain 18 disulfide bonds, with 4 of them being interchain. Table ST1 (in Supporting Information) describes the disulfide linkages. The variants also contain 8 N-glycosylation sites per monomer. Each peptidase domain of the ACE2-Fc variants is capable of binding one SARS-CoV-2 SpFr (Sequence Seq3 in Supporting Information; 183 amino acids), which contains one glycosylation site. The ACE2-Fc/SpFr structure is depicted in Figure 2. Zoomed views of the ACE2/SpFr interface are shown in Figure SF4. The coordinated Zn^2+^ site is shown in Figure SF1. All 3D structures are rendered with VMD (34).

**Figure 2.**
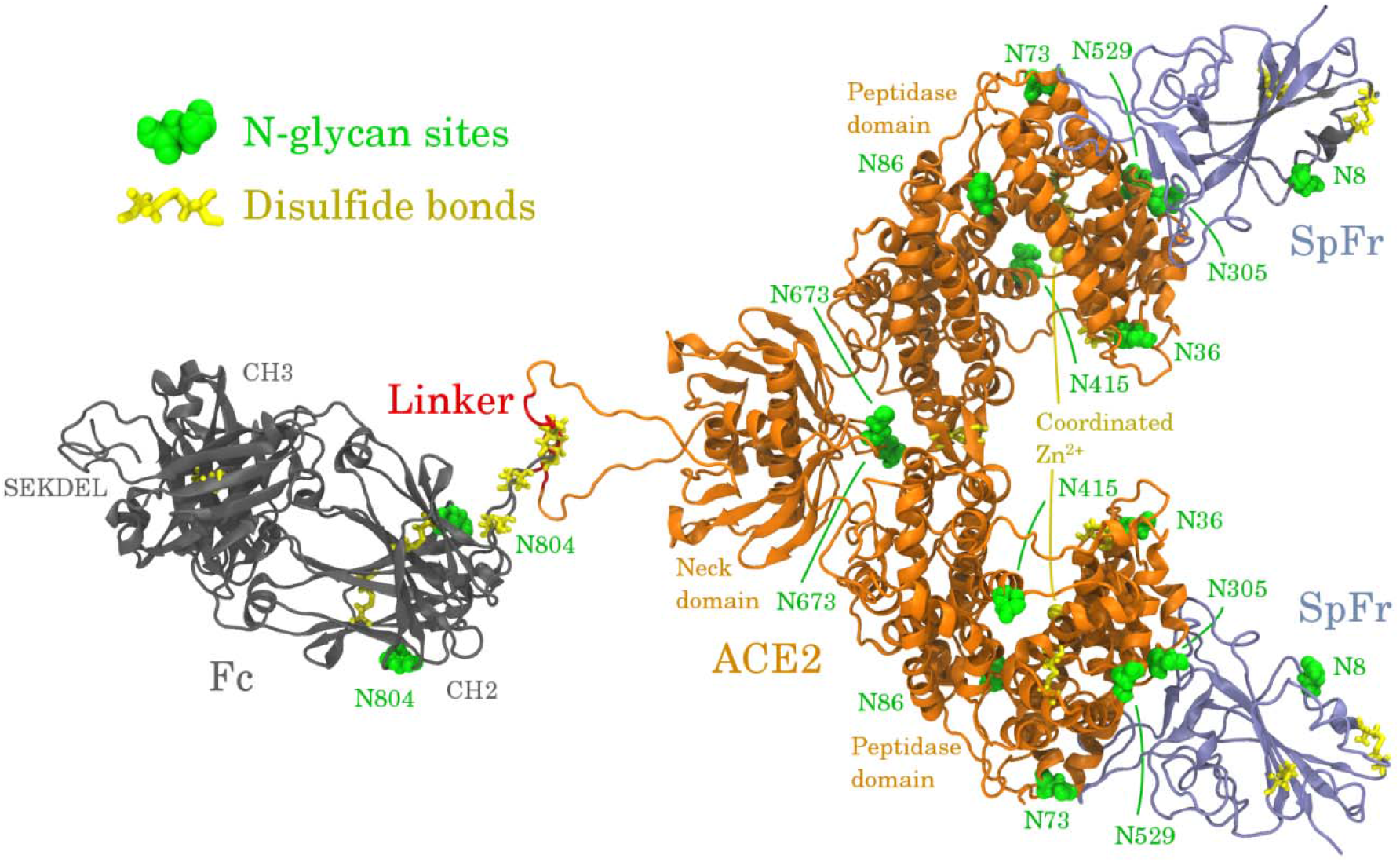
Infographic of the ACE2-Fc variant 1 homodimer bound to two SpFr.

### Simulated systems

ACE2-Fc variant 1 will express high-mannose type glycans when synthesized in plants, while variant 2 will express standard plant glycans. Additionally, SARS-CoV-2 SpFr will exhibit its own glycosylation depending on the host cell; here we assume common mammalian-like glycosylation. Our simulations employed uniform glycosylation profiles to approximate these glycosylation profiles. ACE2-Fc variant 1 is fully glycosylated with Man8 glycans, variant 2 is fully glycosylated with GnGnXF3 glycans the latter is consistent with a recent experimental study (35), and the SpFr is glycosylated with ANaF^6^(36). Figure 3 shows these glycans using the Consortium of Functional Glycomics nomenclature.

**Figure 3.**
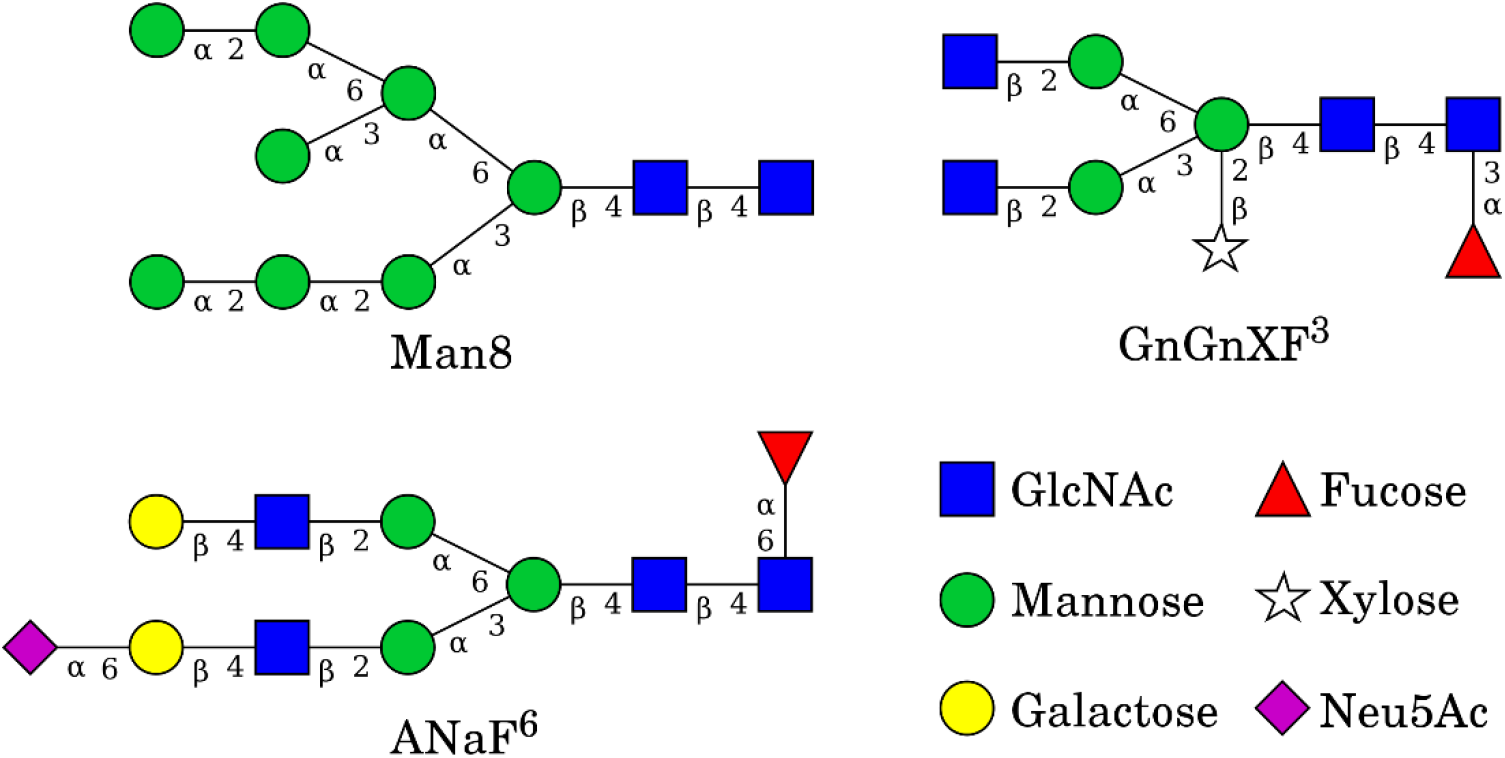
Glycans used in the simulated systems. All structures were built using GlycanBuilder (37).

Four systems containing ACE2-Fc variants were simulated in this work. The first system contains ACE2-Fc variant 1 with Man8 glycans. The second system contains ACE2-Fc variant 2 with GnGnXF^3^ glycans. The third and fourth systems are the immunoadhesins of the first and second systems with the SARS-CoV-2 SpFr bound, respectively. The SpFr is always glycosylated with ANaF^6^. Table 1 summarizes the simulated systems.

**Table 1.**
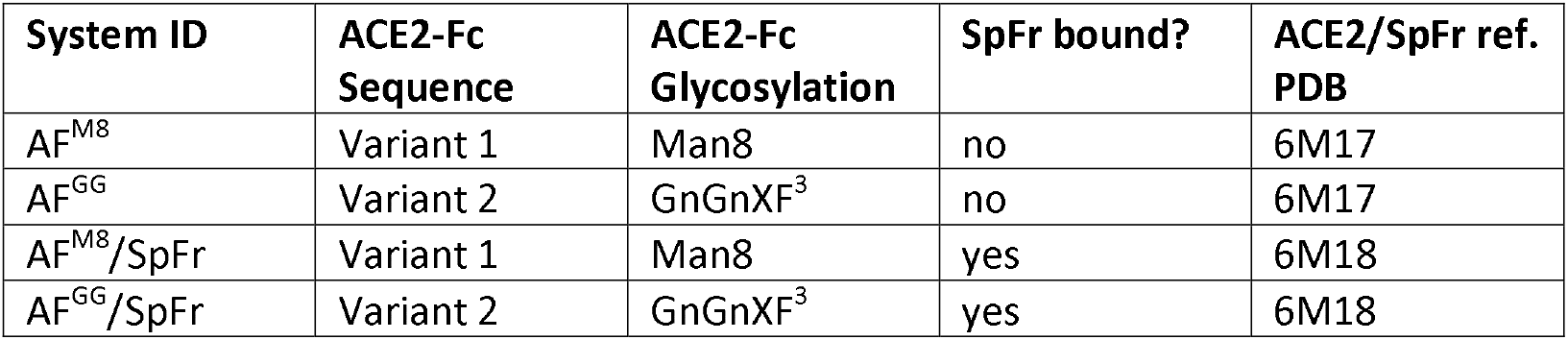
Description of simulated systems.

This work is largely made possible due to the recent cryogenic electron microscopy work that resolves the ACE2-B^0^AT1 and ACE2-B^0^AT1/SpFr structures, corresponding to PDB codes 6M18 and 6M17, respectively (38). The ACE2 and ACE2/SpFr domains were taken from these structures and fused to the Fc domain (PDB 3SGJ) (39). The Zn^2+^ and coordinating residues in 6M17 and 6M18 are poorly coordinated in these structures. The conformation of these residues along with a coordinating water were instead taken from PDB 1R42 (40). Histidine protonation states for each system were determined using Reduce (41), and are summarized in Table ST2.

### Simulation Procedure

The simulation procedure includes the following steps:

1. Fuse ACE2 with Fc to ACE2-Fc using Modeller (42)
2. Model Zn^2+^ and coordinating residues with MCPB.py (43)
3. Attach glycans using glycam.org (44)
4. Merge structures from 2. and 3. using github.com/austenb28/MCPB_Glycam_merge (45)
5. Generate topology files using AmberTools (43)
6. Convert topology files to Gromacs format using Acpype (29, 46)
7. Perform rigid energy minimization (EM) of glycans using github.com/austenb28/GlyRot (47)
8. Perform EM (emtol = 1000 kJ/mol/nm)
9. Solvate and add ions
10. Perform 10 ps constant volume (NVT) (dt = 0.2 fs, T = 310 K)
11. Perform EM (emtol = 1000 kJ/mol/nm)
12. Perform 100 ps NVT (dt = 2 fs, T = 310 K)
13. Perform 100 ps constant pressure (NPT) (dt = 2 fs, T = 310 K, P = 1 atm)
14. Perform 75 ns production NPT (dt = 2 fs, T = 310 K, P = 1 atm)

Steps 2 and 4 are only required for AF^M8^ and AF^M8^/SpFr, since they contain the coordinated Zn^2+^ sites. Steps 4 and 7 exhibit new, publicly available software under GNU General Public Licenses. GlyRot has previously been used to model glycosylated butyrylcholinesterase and CMG2-Fc (13, 14). Forcefield topologies were generated using the AmberFF14SB (48)forcefield for protein atoms, the Glycam06-j (49) forcefield for glycan atoms, and the SPC/E water model (50). Steps 8 through 14 are performed using the Gromacs suite (51–53). Systems were solvated in rectangular boxes such the minimum distance between the solute and periodic boundary is 1.2 nm. A rectangular box (for size see Table ST3) was found to be sufficient for 75 ns; longer simulations may require a larger cubic box if the solute rotates significantly. A reduced timestep NVT in step 10 is required to relax solute-solvent contacts. Steps 10-13 used position restraints on the protein atoms. All simulations were performed at 310K and 1 atm with the Velocity Rescale thermostat (54) and Parinello-Rahman barostat (55) using time constants of 0.1 ps and 2 ps, respectively. All water bonds are constrained with SETTLE (56); all other bonds are constrained with LINCS (57). A 1 nm cutoff was used for short-range nonbonded interactions. Particle Mesh Ewald was used to model long-range electrostatics (58). Table ST3 contains additional information on system sizes and solvation. Each system was simulated using one compute node with 16 cores. Simulations averaged 2.9 ns/day for systems without the SpFr, and 2.0 ns/day for systems with the SpFr.

## Results and Discussion

Figure 3 shows the starting configurations generated as described above (left) and the configurations after MD for 75 ns (right) of all simulated systems. Systems of this size will not fully equilibrate in 75 ns, but evidence of structural stability and concerted motion can still be observed. This is in agreement with a recent equilibration study of a fully glycosylated Spike protein (28). All systems exhibit varying length of the flexible linker domain between ACE2 and Fc during simulation. The domain separation can be quantified by analyzing the center of mass distance between the ACE2 and Fc ordered domains, shown in Figure SF2. AF^M8^ and AF^M8^/SpFr exhibit clearly more shortening of the linker domain than AF^GG^ and AF^GG^/SpFr, possibly due to the difference in glycosylation in the Fc domain, which is closest to the flexible linker region. AF^M8^/SpFr has the shortest distance between the ACE2 and Fc domains after 75 ns, which is consistent with its final configuration shown in Figure 3. In the AF^M8^/SpFr and AF^GG^/SpFr systems, ACE2 glycans near the ACE2/SpFr interface form contacts across the binding interface, indicating that glycosylation may significantly affect binding kinetics. Additionally, glycans on SpFr that were initially oriented away from the protein are reoriented towards ACE2 after 75 ns. The structure of the ordered domains of ACE2-Fc and SpFr appear to retain structural stability. As expected, the glycans, on the other hand, show significant reorientation, as the configurational dynamics of glycans is faster than proteins (13).

**Figure 3.**
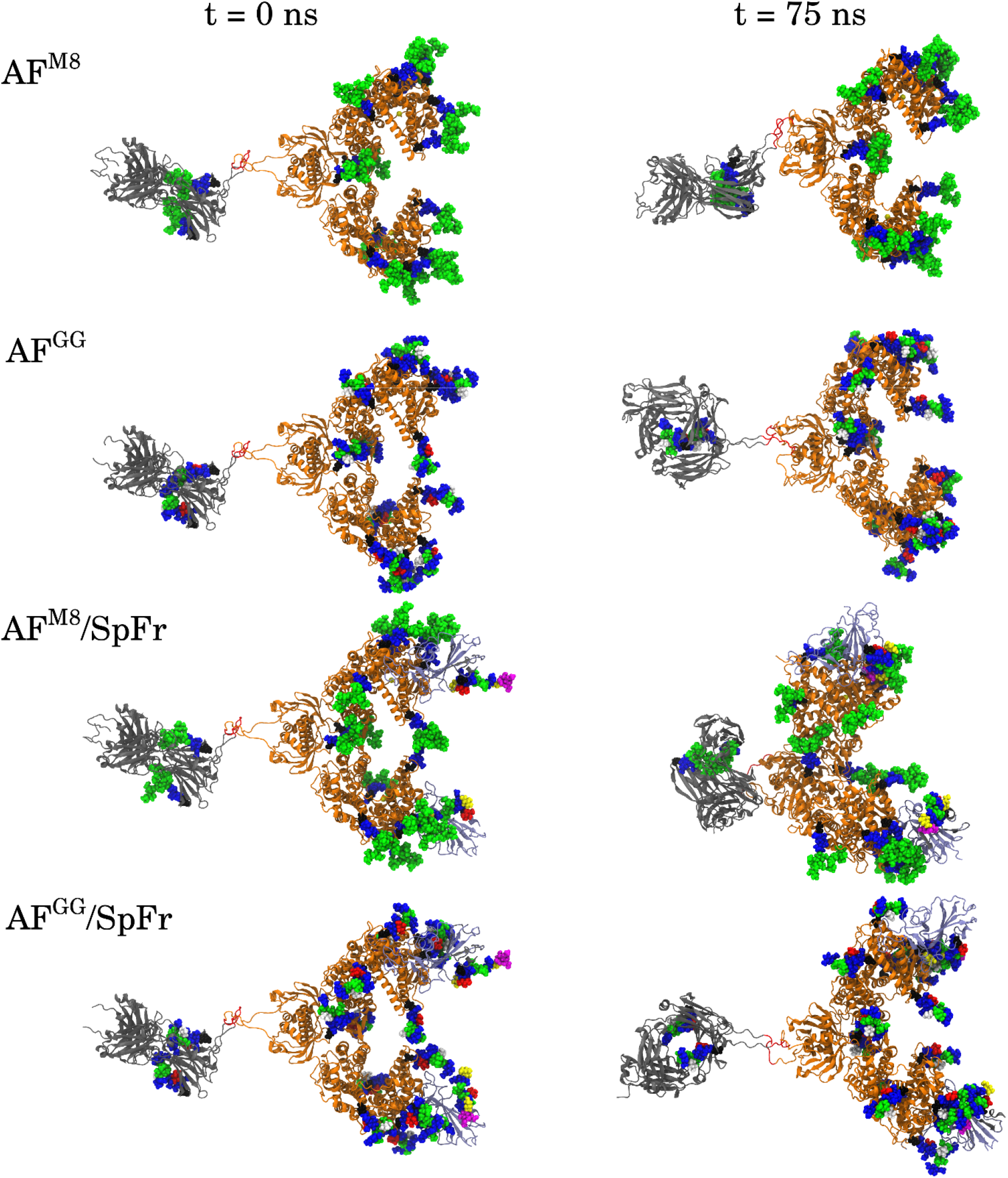
Initial (left) and 75 ns simulated (right) configurations of all systems.

To quantitatively assess structural stability, the root mean square deviation (RMSD) of the ordered domains of ACE2-Fc are shown in Figure 4. All profiles exhibit dynamics near or below 2.5 Å, indicating no major unfolding events have occurred. Conformational trending occurs when the RMSD increases from the initial and decreases towards the final. Conformational trending is evident in the ACE2 domain of all systems. Conformational trending is less evident for the Fc domains, except for the AF^GG^/SpFr system, which exhibits significant conformational trending during the first 20 ns. This difference could indicate that GnGnXF^3^ glycosylation in the Fc domain of the AF^GG^/SpFr promoted a conformational change in the Fc domain. Backbone RMSD profiles for the SpFr are provided in Figure SF3. The SpFr domains show RMSD profiles with significant conformational trending, potentially due to contacts with nearby glycans.

**Figure 4.**
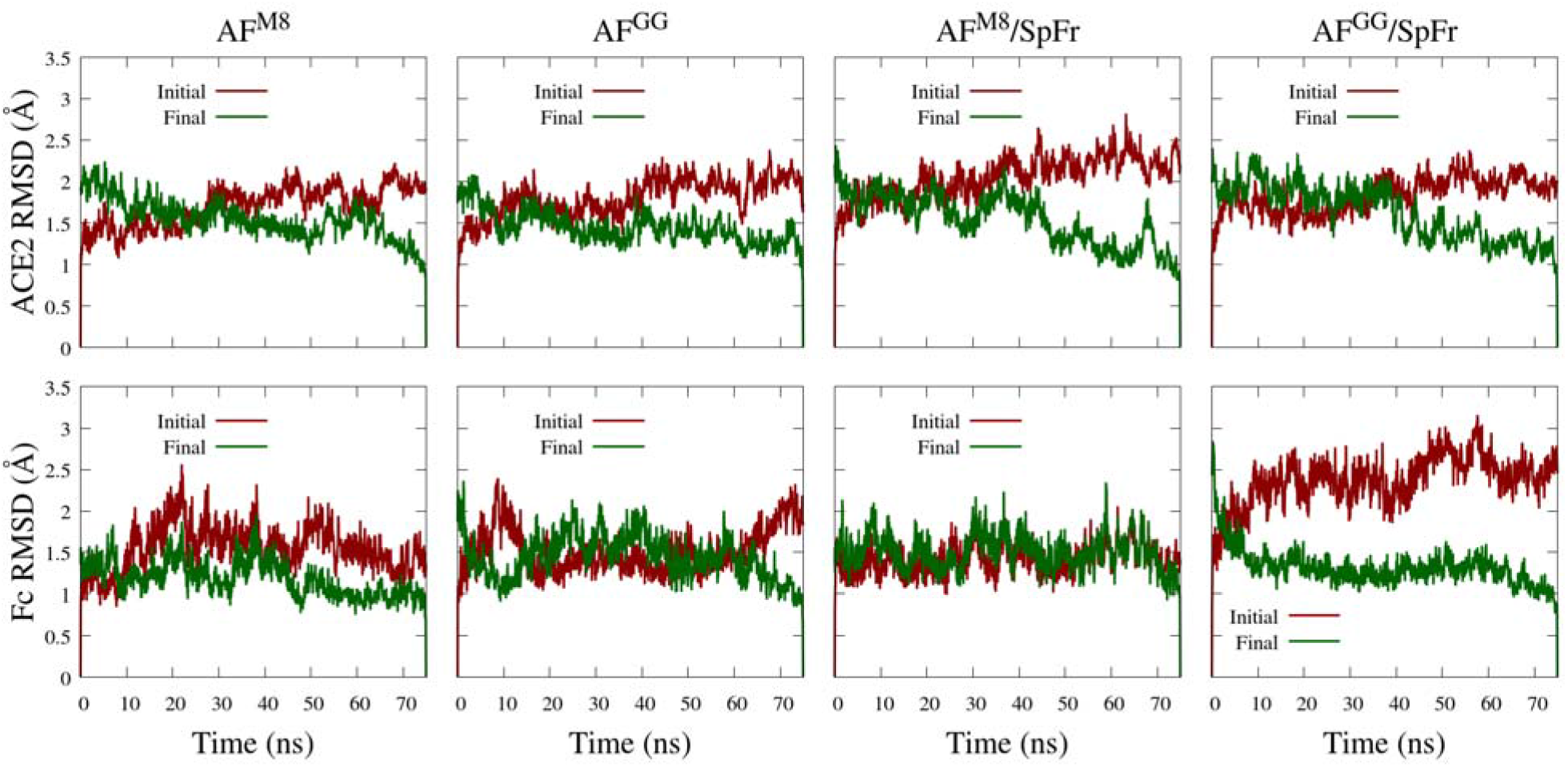
Backbone RMSD profiles of ACE2 (top) and Fc (bottom) ordered domains referenced to initial and final simulation configurations. ACE2: residues 4-707. Fc: residues 745-950. (see SI for sequences)

## Conclusions

We have developed fully glycosylated models of ACE2-Fc immunoadhesins with and without interactions to glycosylated SARS-CoV-2 spike protein fragments. Atomic resolution models can be used to help guide the development of ACE2 and/or ACE2-Fc therapeutics for COVID-19 and potentially other coronavirus borne diseases.

We found that glycosylations affects protein structure, and potentially ACE2/SpFr binding. It is not yet clear how important these differences are, but they must be treated carefully when designing ACE2-Fc variants. The work exhibited here provides a direct avenue for collaborations between experimental and computational researchers.

All models developed here are freely available for researchers and future COVID-19 related simulations. Simulations with a wider variety of glycosylations as well as for longer times are in progress and will be reported in the future. The open-source workflows and tools that have been generated for glycoprotein simulations will be useful for general simulations of glycosylated systems. We hope that glycosylation becomes a standard variable in protein molecular simulations in the near future.

## Acknowledgements

We thank Priya Shah for stimulating discussions. Simulations were performed on the HPC1 computing cluster at UC Davis. This work was partially supported by a COVID-19 Research Accelerator Funding Track project at UC Davis.

## Supporting Information

Supporting information contains all sequence definitions as well as additional analysis. Also, all configurations discussed in this paper are available in pdb format both in supporting information as well as from the datadryad.org (https://doi.org/10.25338/B82G9B). Software tools developed for this project are available on GitHub (45, 47).

