## Supplementary material for "Development and simulation of fully glycosylated molecular models of ACE2-Fc fusion proteins and their interaction with the SARS-CoV-2 spike protein binding domain": SI ResubV.docx

**Variant 1: ACE2WildType(18-740)-SSERKCCVE-IgG1Fc(109-330)- SEKDEL**

NCBI Reference Sequence ID: NP_001358344.1 (ACE2); UniProtKB Sequence ID: P01857 (IgG1Fc_human corresponds to amino acid residues 109 - 330)

PDB codes: 6M17 or 6M18 (ACE2), 1R42 (Zn + coordinating residues + coordinating water), 3SGJ (Fc)

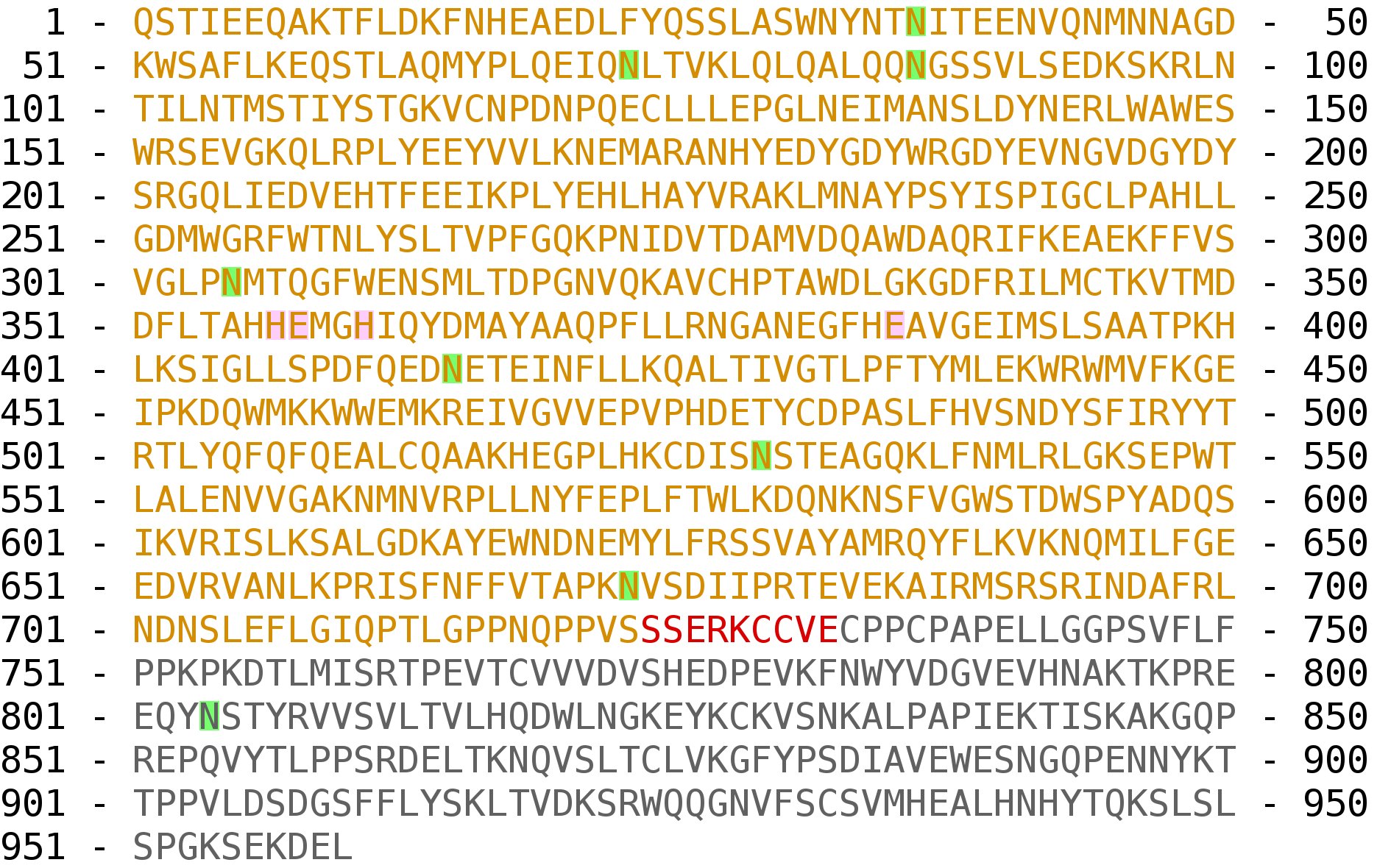

Sequence Seq1. ACE2-Fc variant 1 sequence. ACE2 in orange, linker in red, Fc in grey. Glycosylation sites highlighted in green. Coordinating Zn^2+^ residues highlighted in pink.

**Variant 2: ACE2Mutant(18-740,H374N,H378N)-SSERKCCVE-IgG1Fc(109-330)**

NCBI Reference Sequence ID: NP_001358344.1 (ACE2), UniProtKB Sequence ID: P01857 (IgG1Fc_human corresponds to amino acid residues 109 – 330)

PDB codes: 6M17 or 6M18 (ACE2), 3SGJ (Fc)

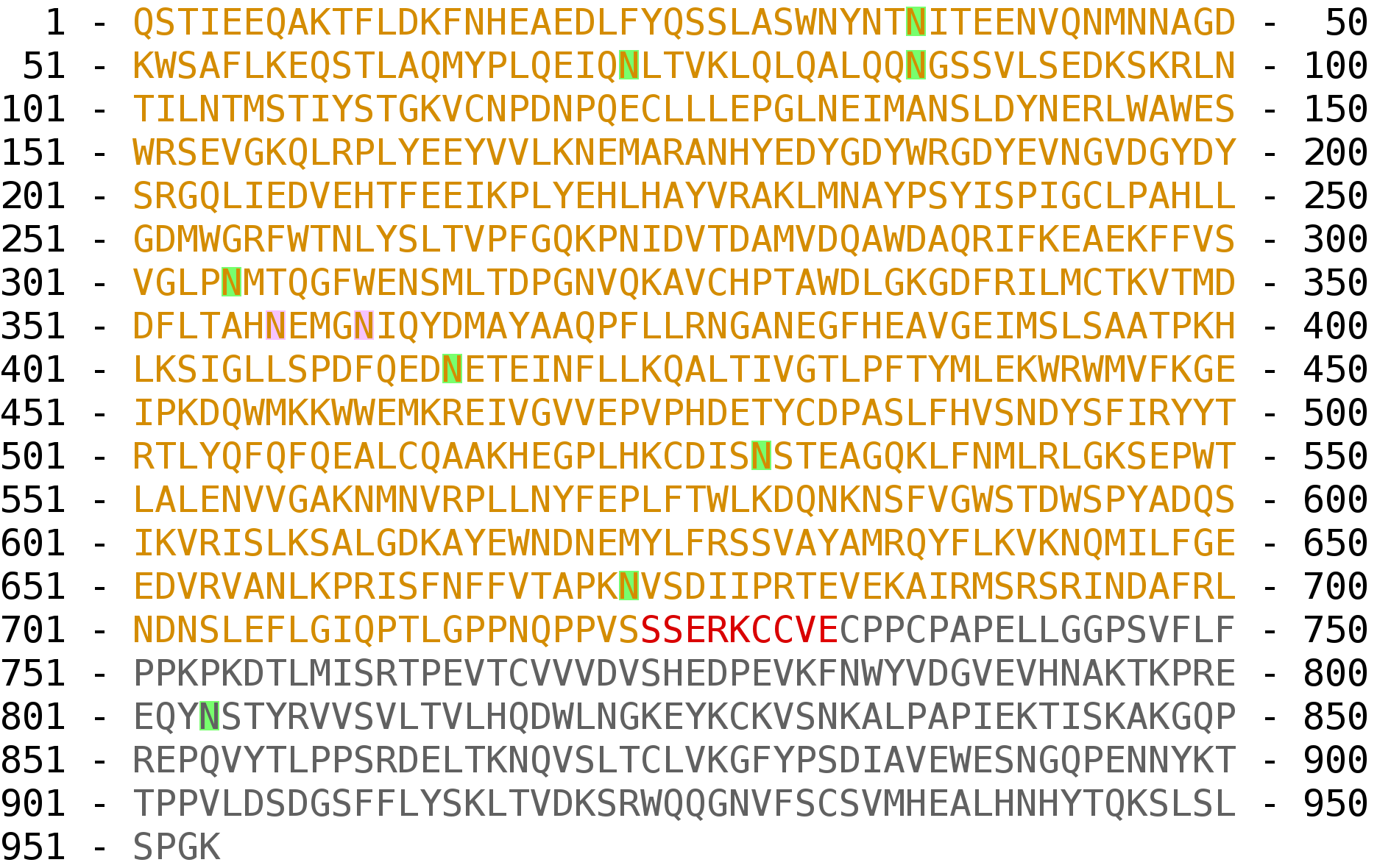

Sequence Seq2. ACE2-Fc variant 2 sequence. ACE2 in orange, linker in red, Fc in grey. Glycosylation sites highlighted in green. Mutated residues highlighted in pink.

**SpFr (crystallized residues only):**

NCBI Reference Sequence ID: YP_009724390.1(SpFr corresponds to amino acid residues 336 – 518)

PDB codes: 6M17

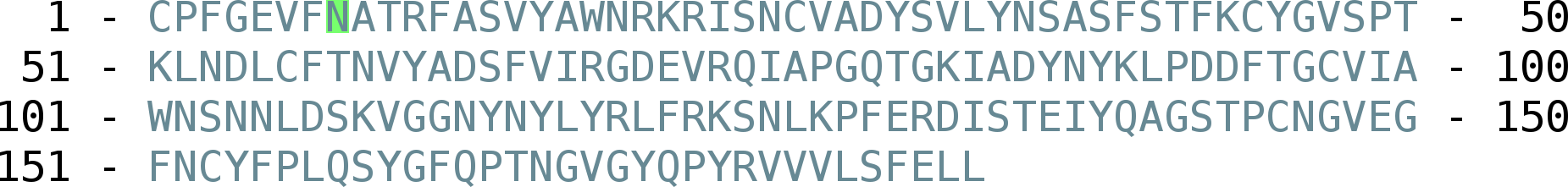

Sequence Seq3. Spike fragment (SpFr) sequence. Glycosylation site highlighted in green.

Table ST1. Table of disulfide bonds. Interchain disulfide bonds are specified using A and B.

| **ACE2-Fc** | | | | **SF** | |
| --- | --- | --- | --- | --- | --- |
| CYS 1 | CYS 2 | CYS 1 | CYS 2 | CYS 1 | CYS 2 |
| 116 | 124 | 733A | 733B | 1 | 26 |
| 327 | 344 | 736A | 736B | 44 | 97 |
| 513 | 525 | 768 | 828 |  |  |
| 729A | 729B | 874 | 932 |  |  |
| 730A | 730B |  |  |  |  |

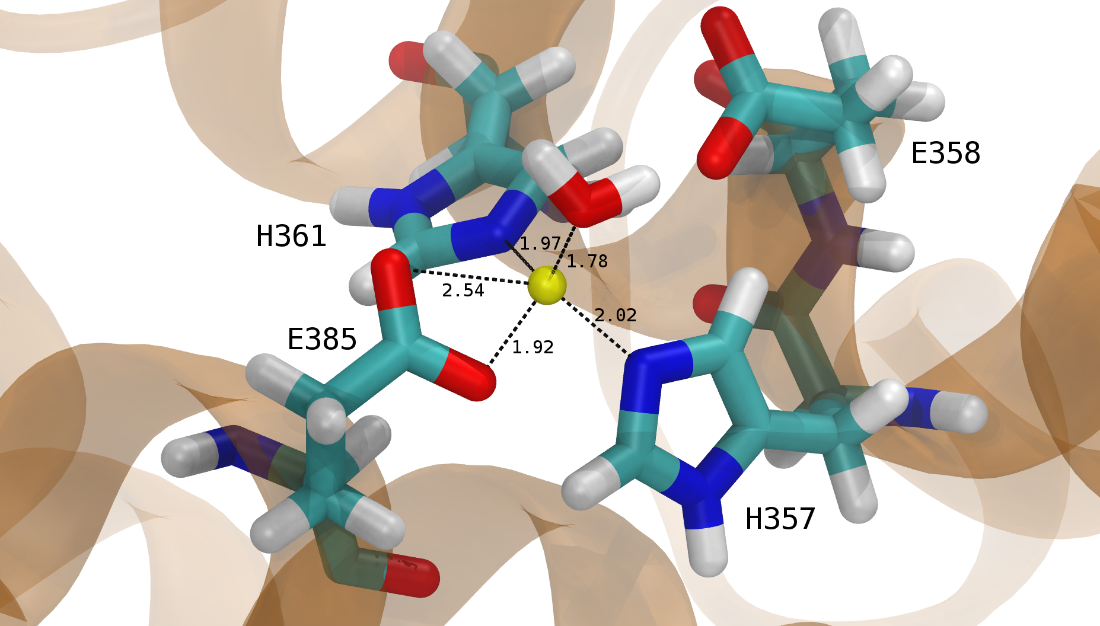

Figure SF1. Coordinating residues of the Zn^2+^ (yellow) active site of variant 1. Coordinating bonds with their distances in angstroms are labeled.

Table ST2. List of delta-protonated histidines for simulated systems. All other histidines are epsilon-protonated. Indices are for ACE2-Fc; SpFr has no histidines.

| AF^M8^ | AF^GG^ | AF^M8^/SpFr | AF^GG^/SpFr |
| --- | --- | --- | --- |
| H17 | H17 | H356 | H356 |
| H357 | H942 | H357 | H942 |
| H361 |  | H361 |  |
| H942 |  | H942 |  |

Table ST3. Additional simulation details.

| System ID | Initial Box dimensions  (nm x nm x nm) | # waters | # Na^+^ | # Cl^-^ |
| --- | --- | --- | --- | --- |
| AF^M8^ | 18.0 x 15.5 x 22.0 | 188975 | 619 | 573 |
| AF^GG^ | 17.6 x 16.3 x 22.7 | 197092 | 644 | 598 |
| AF^M8^/SF | 18.0 x 16.7 x 24.2 | 222602 | 721 | 677 |
| AF^GG^/SF | 17.9 x 16.7 x 24.2 | 221872 | 718 | 674 |

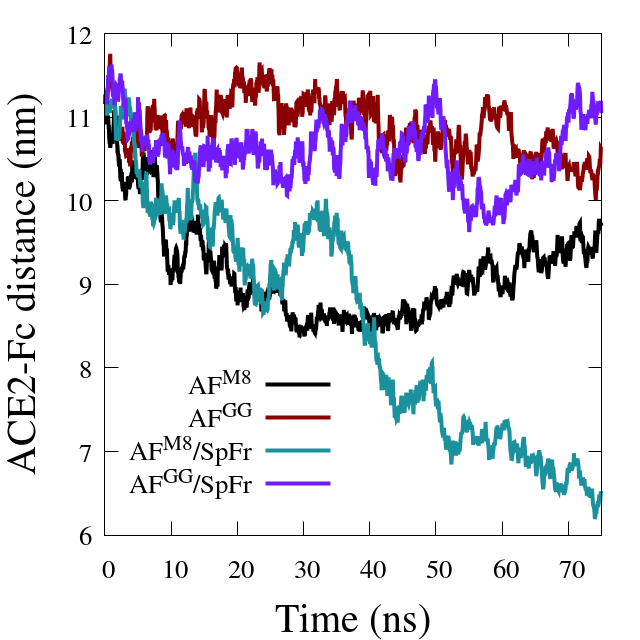

Figure SF2. Center of mass distance between the ordered domains of ACE2 and Fc. ACE2: residues 4-707. Fc: residues 745-950.

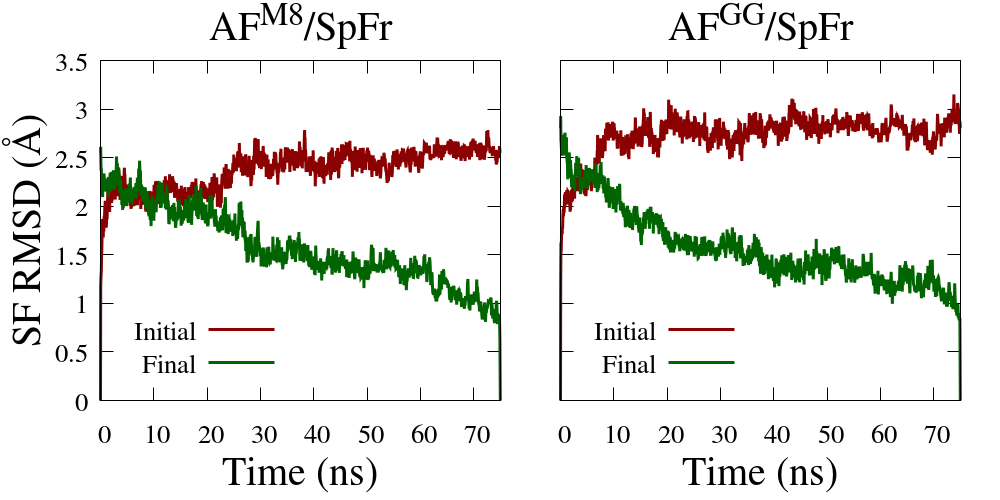

Figure SF3. Backbone RMSD profiles of the SpFr referenced from initial and final simulation configurations.

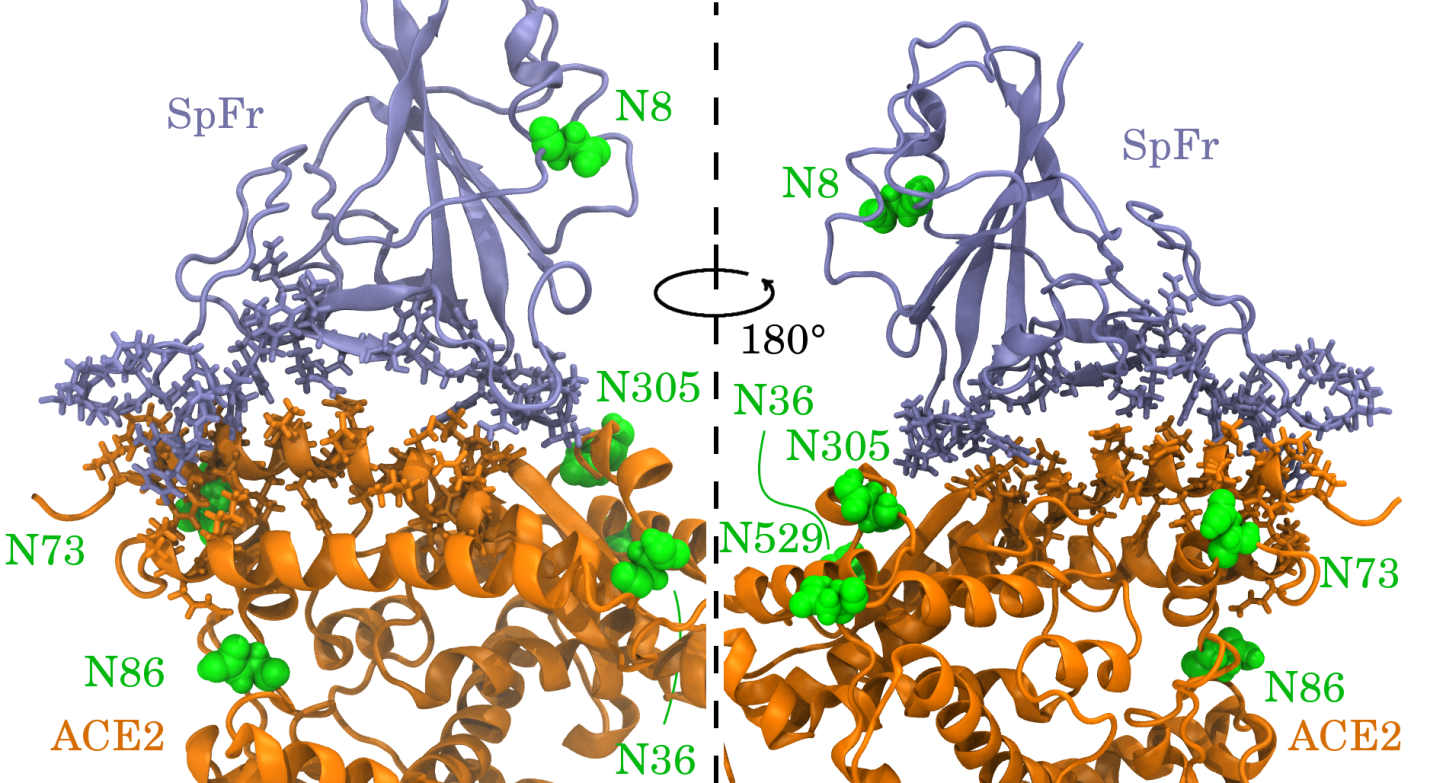

Figure SF4. Front (left) and back (right) zoom views of the interface between the ACE2 and SpFr domains. Interfacial residues are shown in licorice.
